# Protein Multiple Alignments: Sequence-based vs Structure-based Programs

**DOI:** 10.1101/413369

**Authors:** Mathilde Carpentier, Jacques Chomilier

## Abstract

Facing the huge increase of information about proteins, classification has reached the level of a compulsory task, essential for assigning a function to a given sequence, by means of comparison to existing data. Multiple sequence alignment programs have been proven to be very useful and they have already been evaluated. In this paper we wished to evaluate the added value provided by taking into account structures. We compared the multiple alignments resulting from 24 programs, either based on sequence, structure, or both, to reference alignments deposited in five databases. Reference databases, on their side, can be split in two: more automatic ones, and more manually ones. Scores have been attributed to each program. As a global rule of thumb, five groups of methods emerge, with the lead to two of the structure-based programs. This advantage is increased at low levels of sequence identity among aligned proteins, or for residues in regular secondary structures or buried. Concerning gap management, sequence-based programs place less gaps than structure-based programs. Concerning the databases, the alignments from the manually built databases are the more challenging for the programs.

## INTRODUCTION

Multiple alignments of protein sequences constitute an essential tool to explore evolution, diversity, conservation and function of proteins (1–4). Despite the impressive increasing number of available structures, most of these alignments are still computed by software relying only on sequence information. Protein structures are mostly used in a second step to manually refine the alignment (5). Since it is generally admitted that structures are more conserved than sequences (6), structural information may guide a particularly difficult alignment of very divergent proteins (7). Nevertheless, multiple protein structure alignment methods, or methods combining sequence and structure, are not widespread.

A structural alignment may outline other types of information than homology (8). In protein sequence alignments we align amino acids which are considered as homologous, i.e. deriving from an ancestral sequence by substitution at the same site. In structural alignments, we align positions similar from the point of view of local and/or global conformations. This structural similarity does not always imply homology (8). Indeed, sub domain fragments can be found in many different folds, with unrelated functions or various origins (9–11). The conceptual model behind sequence alignment explicitly considers three events for evolution: insertion, deletion and mutation. The model behind structure alignment is not so clear, partly because of the folding step of protein structures.

From all the previous arguments, it is difficult to claim that structure alignments provide the golden standard to evaluate the quality of sequence alignment. This is particularly the case when the proteins have a low level of similarity or if the homology of the whole genes is questionable. However, as structures are better conserved, alignments should be more reliable when information from sequences and structures are combined. We therefore compared the alignments computed from structure or both structure and sequence with those from sequence only.

Multiple sequence alignment methods have been compared in many articles and with several types of benchmarks reviewed in (12). The most widely used benchmarks are composed of a collection of reference alignments considered as the gold standard. The reference alignments are constructed mainly from the sequence and structural information, but also according to other information as the function. Some of them are manually curated. When a new alignment method or an improvement is published, its performance is usually assessed by comparison to other methods by aligning the proteins of these reference alignment databases. There are also many comparative studies of the performances of sequence alignment methods (13, 14). The second type of benchmarks relies on simulated sequences (15). A third type of benchmarks relies on a direct comparison of all computed alignments, without any reference alignment (16, 17). The fourth type of benchmarks is to calculate phylogenetic trees from the alignments and to check their validity (18). For structure-based alignment methods, less comparative studies of have been conducted and most of them compare pairwise structural alignment programs (8, 19–24). Multiple structural alignment programs are compared in the study of Berbalk et. al. (25). The authors first remarked that the programs were generally very difficult to use and that there is room for improvements concerning usability and applicability. They concluded that combining different alignment approaches into a single program supported by an automated scoring could improve the alignment quality but that until such a method is implemented, it seems important for a user to apply different tools and to manually compare their results.

We are not aware of a thorough comparative study of the performance of sequence-based and structure-based programs. We believe that such a study is important to address some questions: Are structure-based methods really superior to retrieve homologous residues? Or is it the sequence and structure ones? In what cases should we use structure methods, sequence and structure or sequence-based methods? These are the aims of this article.

## MATERIAL AND METHODS

### Databases

In this study, we used the most widely used type of benchmarks: the reference multiple alignments built from sequences, structures and function information, and considered as the gold standard. The usage of simulated sequence is not possible in our case because there is no structure associated. It is possible to compare all alignments without a reference but as programs may be consistently wrong; therefore we decided to avoid this approach in this article. The phylogeny-based approach would be very interesting but it requires a database of validated trees of genes with all known protein structures, which is beyond the scope of the article.

We have selected 846 alignments, containing at least three protein chains, from five reference multiple alignment databases: BALIBASE 2 (26), BALIBASE 3 (27), HOMSTRAD (28), OXBENCH (29) and SISYPHUS (30). Some alignments have been discarded: those with two or more proteins with identical amino acid sequence, NMR or theoretical model structures, structures with missing residues and those with various inconsistencies. We did not consider the alignments of other well-known databases listed in (31) for various reasons: PREFAB (32) because it is composed of pairwise alignments; IRMbase (33) because there is no structure associated to the simulated fragments and SABMARK (34) because of some inconsistencies in the multiple alignments which are built from pairwise structural alignments, pointed by the author and in (35). We also had difficulties accessing PALI (36) and couldn’t download the database. From all the databases, we only consider the core of the alignments but its definition depends on the database.

We have selected 29 families from BALIBASE 2 (BB2) and 38 from BALIBASE 3 (BB3), manually curated by checking the alignments of functional and other conserved residues. In each family, all proteins share the same structural fold, so the core can be reliably defined, excluding ambiguous or non-superimposable regions, unrelated secondary structure borders or some loop regions. HOMSTRAD, from which we selected 357 families, is exclusively based on proteins with known structures, and each family is aligned with the programs MNYFIT (37), STAMP (38) and COMPARER (39). These produce structure-based alignments that are annotated with JOY (40) and individually examined and modified if necessary. JOY produces core blocks annotations defined as the regular secondary structures elements. We retrieved from OXBENCH 330 alignments from the subset multi with three or more proteins in each, not split in domains (full-length sequences). The multiple alignments are computed by STAMP (38). All the aligned positions were taken as the core blocks. The last database, SISYPHUS, is based on the families of domains from the structural classification SCOP (41) with non-trivial structural relationships. Multiple alignments are manually constructed for structural regions that range from oligomeric biological units, or individual domains to fragments of different size and are manually curated. Sisyphus annotates the structurally equivalent residues in the alignments and we consider them as the core blocks.

Many structure-based alignment programs don’t output all the residues of input protein structures (some residues are removed or ignored) or change the name of the sequences. We have developed two programs to solve this issue: the first one retrieves the correspondence of the protein between the reference alignment and the output alignment and the second makes each sequence of an output alignment identical to the sequence in the reference alignment.

### Alignment Quality Evaluation

The alignments produced by each program are evaluated by comparison with the reference alignments by two scores, following Thomson et al (13): 1) the fraction of pairs of residues in the reference alignment correctly identified by a given method, known as the Sum-of-Pairs (SP) score; 2) the Column Score (CS) score, which describes the fraction of reference columns identified. As usually done in alignment method comparisons (13, 42, 43), Friedman tests (44) were performed. This test is more conservative than the Wilcoxon test that assumes a symmetrical difference, and this is not always the case. All tests, plots and heatmaps have been done with R (45). The average multiple RMS have been computed with THESEUS (46) that has been applied to all alignments, reference ones or computed by the tested programs. We have counted the number of gaps in all columns between the first and last core elements. We present in the article only the proportion of columns containing one or more gap opening. Solvent accessibility (ASA) is calculated with NACCESS for all the proteins, in order to separate the amino acids in two classes: either buried (relative ASA < 25%) or exposed (47). Secondary structure assignments have been performed with STRIDE (48). The six classes given in the output of STRIDE are back coded in three classes: helices, strands and coils. All analyses have been made according to these characteristics: the residues of the core blocks have been attributed either as buried or accessible, and either in helix, strand or other (loop).

### Programs

We have 3 categories of multiple alignment programs: sequence-based, sequence+structure-based and structure-based. We only included programs respecting the following conditions: (i) available for download, (ii) output a file containing the alignment in a standard format, (iii) run without error. Each multiple alignment had to be computed in less than two hours. Some programs failed to produce enough alignments to allow a significant analysis of their performance and were excluded if the produced an alignment for less than 70% of the dataset. As we mainly aim at addressing the performance of structure-based or sequence+structure-based alignment methods, we tried to be as exhaustive as possible for them. We searched or tested more than 40 programs but many were unavailable or didn’t respect our criteria. We were also surprised by the few number of sequence+structure alignment methods. We didn’t include methods improving alignments afterwards like STACCATO (49). There is a great number of sequence-based programs and we only tested the most popular according to the last studies (14, 50). All the programs included in our study are listed with a short description in Table 1. We have selected 9 sequence-based programs, 5 sequence+structure-based programs, (TCOFFEE/3DCOFFEE is either run with SAP or TM-ALIGN) and 10 structure-based programs.

**Table 1:**
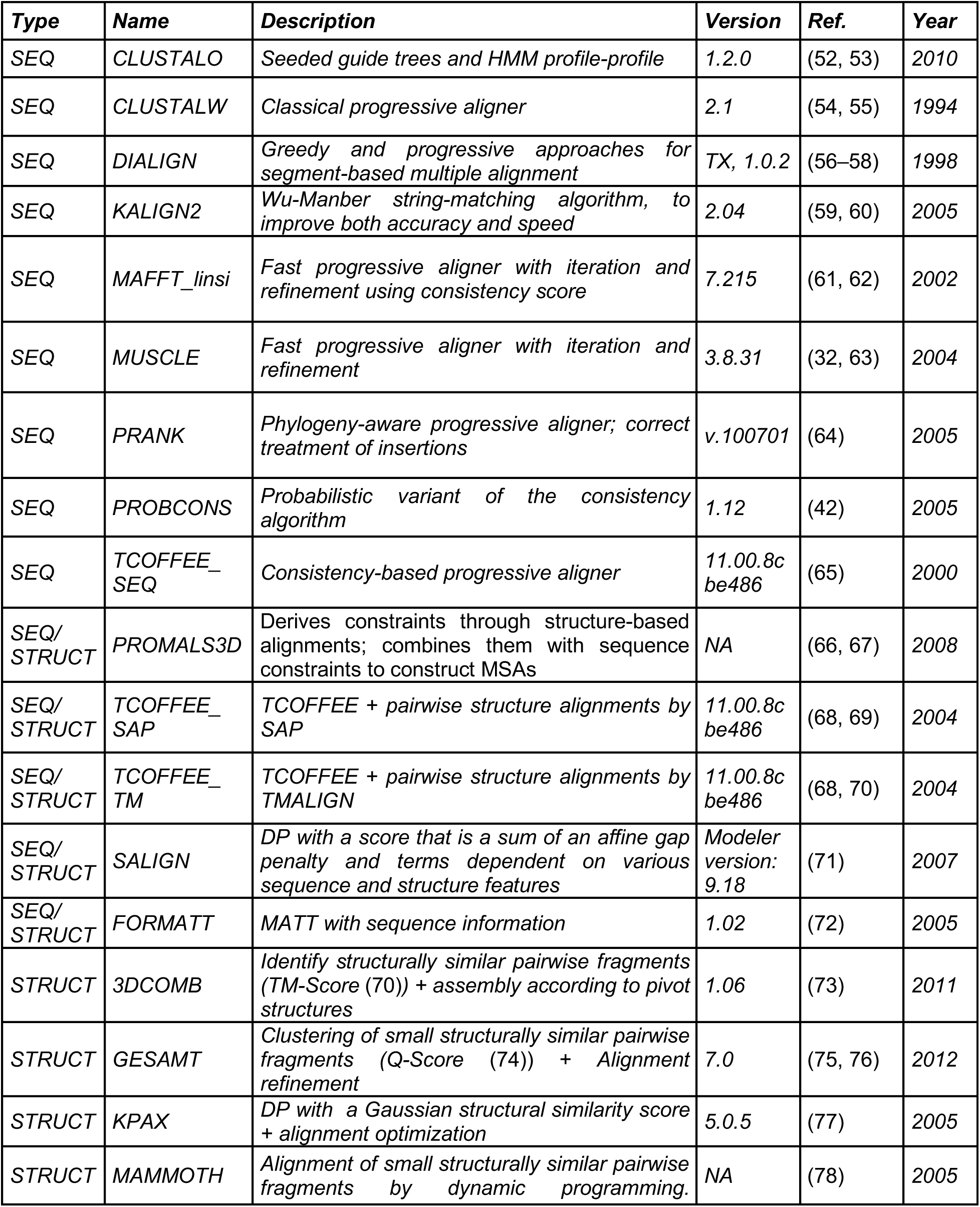
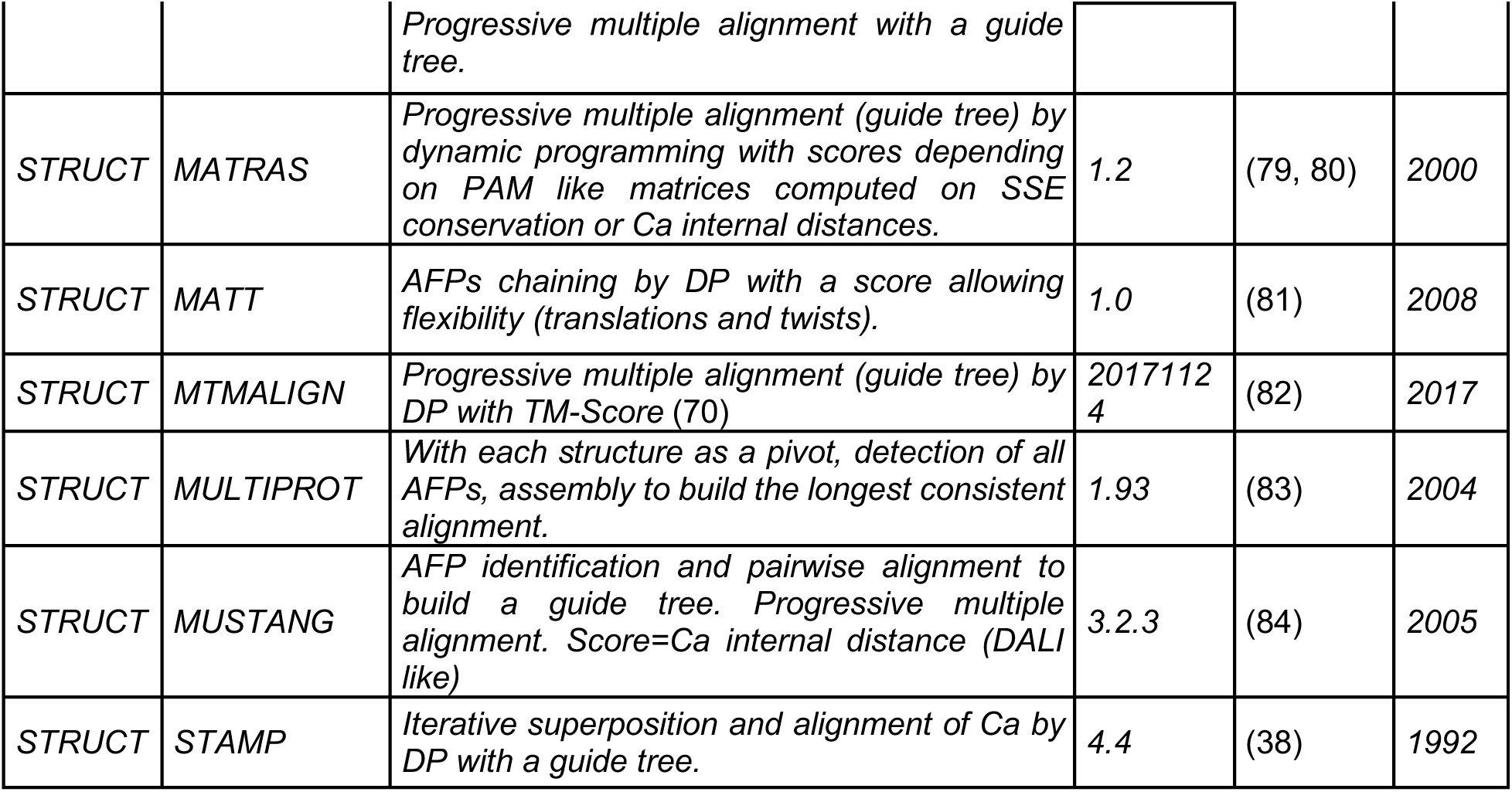
Programs used in this study to align families of proteins from the reference datasets. Categories of programs: SEQ is a sequence-based alignment method; STRUCT is a structure-based alignment method; SEQ/STRUCT is a sequence+structure based program. DP=Dynamic Programming, AFP=Aligned Fragment Pairs, SSE=secondary structure element.

## Results

### Number of computed alignments

All programs have been run on the 846 alignments. All the sequence-based programs were able to calculate all the alignments but some programs of the two other categories failed for some alignments. The proportion of successful alignments is reported in Table 2. Only MATRAS, TCOFFEE_SAP, TCOFFEE_TM, and KPAX, successfully computed all alignments. The failure causes were sometimes the time limit, but most of the time, the programs returned some errors. MAMMOTH encountered the most failures; there is obviously a limit of 25 proteins per alignment for it. The greatest number of failures for all programs is with the SISYPHUS database which is not surprising because it is a benchmark built to be challenging. To improve the robustness of our analysis, we decided to restrict our analysis to the alignments computed by all programs, resulting in 531 alignments: 24 from BB2, 24 from BB3, 288 from HOMSTRAD, 155 from OXBENCH and 40 from SISYPHUS. These 531 alignments involve 2043 chains.

**Table 2:** Number of computed alignments from structure-based or sequence+structure methods.

|  | <u>ALL</u> | <u>BB2</u> | <u>BB3</u> | <u>OXBENCH</u> | <u>HOMSTRAD</u> | <u>SISYPHUS</u> |
| --- | --- | --- | --- | --- | --- | --- |
| MATRAS | 846 100,0% | 29 100,0% | 38 100,0% | 330 100,0% | 357 100,0% | 93 100,0% |
| TCOFFEE(SAP/TM) | 846 100,0% | 29 100,0% | 38 100,0% | 330 100,0% | 357 100,0% | 93 100,0% |
| KPAX | 845 100,0% | 29 100,0% | 38 100,0% | 330 100,0% | 357 100,0% | 93 100,0% |
| PROMALS3D | 845 99,9% | 29 100,0% | 38 100,0% | 330 100,0% | 357 100,0% | 92 98,9% |
| GESAMT | 840 99,3% | 28 96,6% | 38 100,0% | 330 100,0% | 352 98,6% | 93 100,0% |
| 3DCOMB | 838 99,1% | 29 100,0% | 38 100,0% | 326 98,8% | 356 99,7% | 90 96,8% |
| MTMALIGN | 836 98,8% | 29 100,0% | 38 100,0% | 330 100,0% | 356 99,7% | 84 90,3% |
| FORMATT | 831 98,2% | 29 100,0% | 38 100,0% | 328 99,4% | 354 99,2% | 83 89,2% |
| STAMP | 825 97,5% | 29 100,0% | 38 100,0% | 330 100,0% | 357 100,0% | 72 77,4% |
| MATT | 823 97,3% | 29 100,0% | 38 100,0% | 322 97,6% | 354 99,2% | 81 87,1% |
| MUSTANG | 813 96,1% | 29 100,0% | 38 100,0% | 326 98,8% | 350 98,0% | 71 76,3% |
| SALIGN | 802 94,8% | 29 100,0% | 37 97,4% | 318 96,4% | 345 96,6% | 74 79,6% |
| MULTIPROT | 761 90,0% | 28 96,6% | 38 100,0% | 255 77,3% | 357 100,0% | 84 90,3% |
| MAMMOTH | <u>621 73,4%</u> | <u>24 82,8%</u> | <u>24 63,2%</u> | <u>212 64,2%</u> | <u>306 85,7%</u> | <u>56 60,2%</u> |
| #Alignments | 846 | 29 | 38 | 330 | 357 | 93 |

### Databases

The distribution of mean pairwise sequence identity among the 531 multiple alignments of the databases is given in Figure 1. BB2, BB3 and SISYPHUS databases are more focused on low identity, while HOMSTRAD and OXBENCH present alignments of high level of identity.

The proportion of amino acids included in regular secondary structures in the complete dataset is 60%; but, restricted to the core alignments, the proportion increases to 79%.

We checked if some protein families were present in several datasets. We found some chains in several databases even if all the proteins of the family are not the same. The number and proportion of chains included in two databases are listed in Supplementary Table 1. There is some overlap between BB2 and BB3: 48 chains are present both in BB2 and BB3. However, the protein families are different between BB2 and BB3 so we decided to keep them all. The overlaps are very weak for the other datasets.

**Figure 1:**
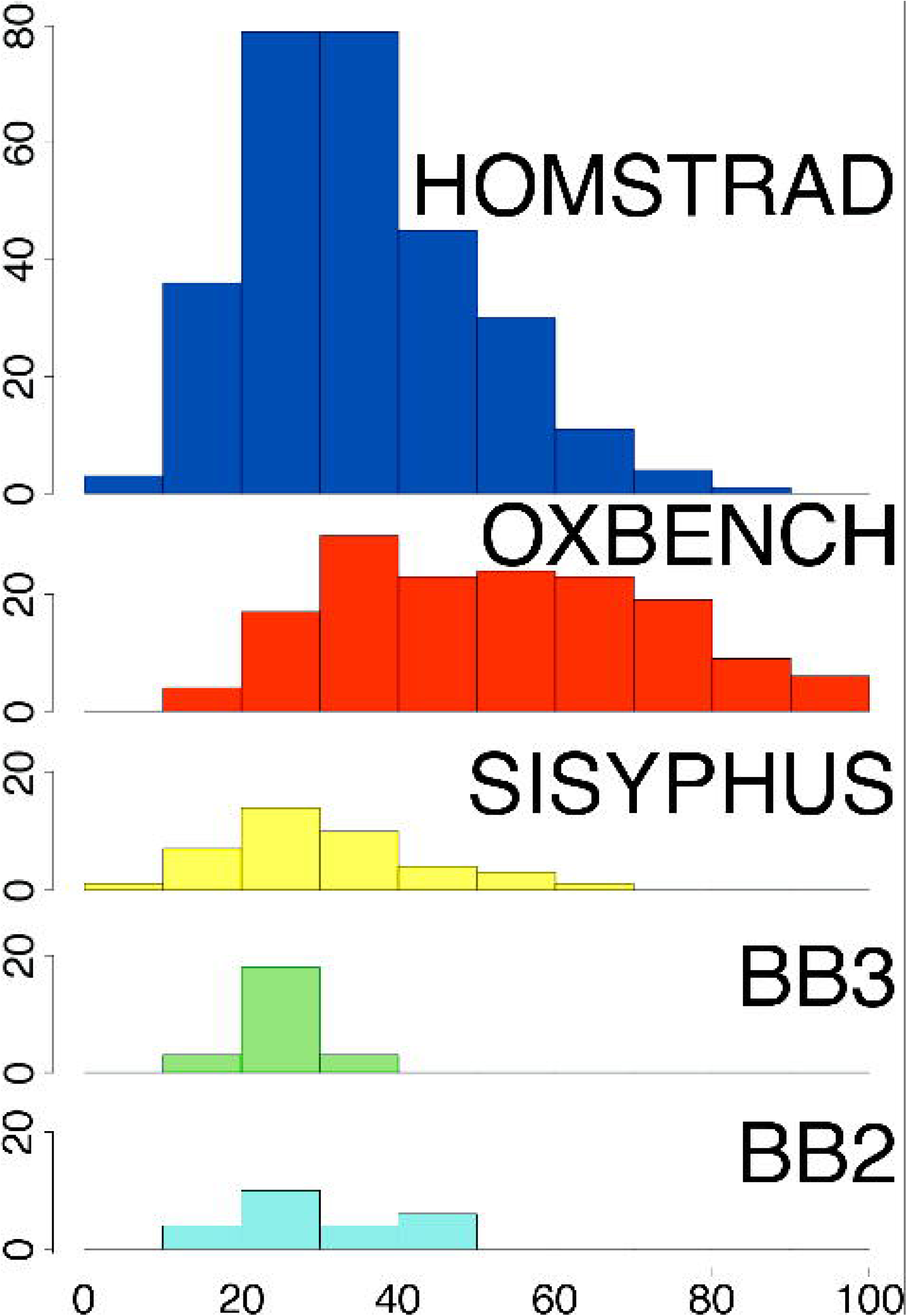
Distribution of core block sequence identity percentage among the five databases. X-axis: identity percentage, Y-axis: number of reference alignments.

### Global Analysis of Alignment Scores

The boxplot distribution of SP and CS scores of each program run on the 531 alignments are presented in Figure 2. The exact median values are reported in Table 2 of the Supplementary Data. Globally, for all programs, the results are impressively good: the SP score medians range from 0.86 to 0.97, meaning that in half of the alignments, more than 86% of the residue pairs are correctly aligned by any methods. Similarly, in half of the alignments, more than 81% of the alignment columns are correct. The scores vary with the programs and it is strikingly clear that structure-based alignment programs have globally better results, except for MULTIPROT. The sorting is roughly the same for SP and CS scores except for FORMATT and MULTIPROT. STAMP has the greatest variability in its results and it is not the best despite the fact that it has been used to build the alignments of two databases (HOMSTRAD and OXBENCH). It is interesting to notice that FORMATT, a modified version of MATT to include sequence information, achieves worse than MATT. It highlights the difficulty to combine sequence and structure information. It is nevertheless possible: TCOFFEE_TM is the best sequence+structure program and it achieves clearly better than TCOFFEE_SEQ. It is however surprising that sequence+structure based methods do not achieve better than structure only methods: their alignments rely on the same structural information, enhanced with sequence information.

**Figure 2:**
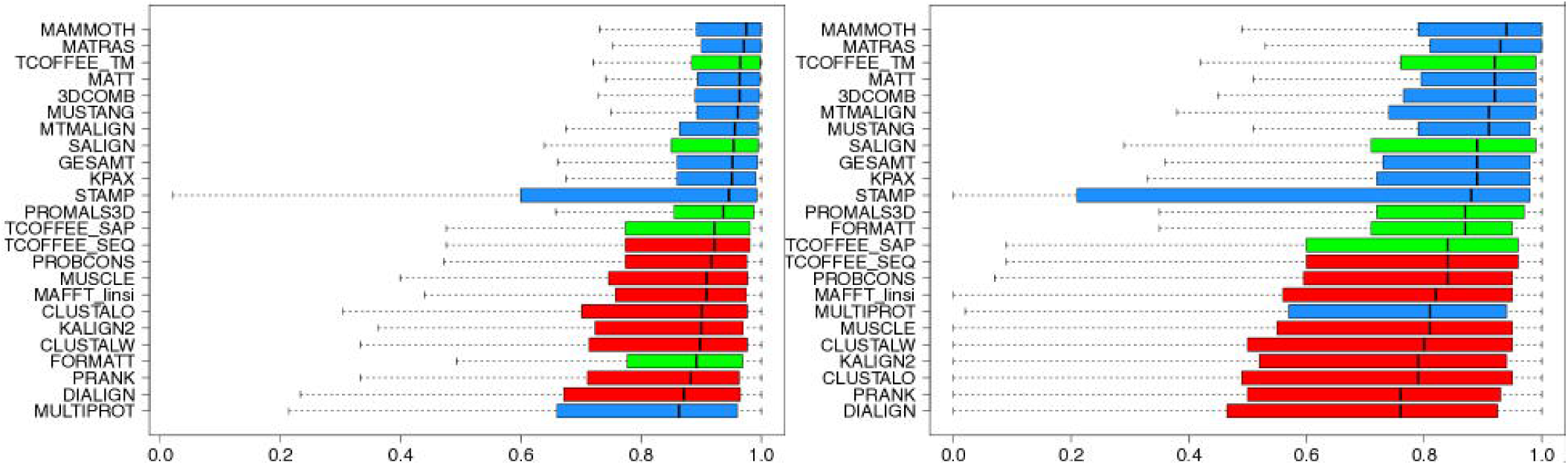
SP (left) and CS (right) Scores: 531 alignments computed by all methods. Programs are sorted according to the median. The colors of the boxes are: red for sequence-based alignment programs, blue for structure-based alignment programs, green for sequence+structure based alignment programs.

For each pair of programs, the significance of their differences has been evaluated by a Friedman rank test on their scores calculated for all 531 alignments (see method). In Figure 3, five groups of methods appear: the differences are mostly non-significant between the programs within each group, but they are significant with the programs outside the groups. The two first groups (blue symbols on the diagonal) contain all structure-based alignment methods but MULTIPROT and three of the sequence+structure-based methods. The next three groups (red symbols) contain all sequence-based alignment methods and two sequence+structure-based methods. The results of TCOFFEE_SAP are identical to TCOFFEE_SEQ. STAMP is particular: its results greatly vary and it is not significantly better or worse than the programs in the two middle groups. MULTIPROT is also particular: the differences are not significant for the SP scores with DIALIGN, PRANK, CLUSTALW and KALIGN2, but for the CS scores, the differences are not significant with all sequence-based methods but PRANK and DIALIGN.

**Figure 3.**
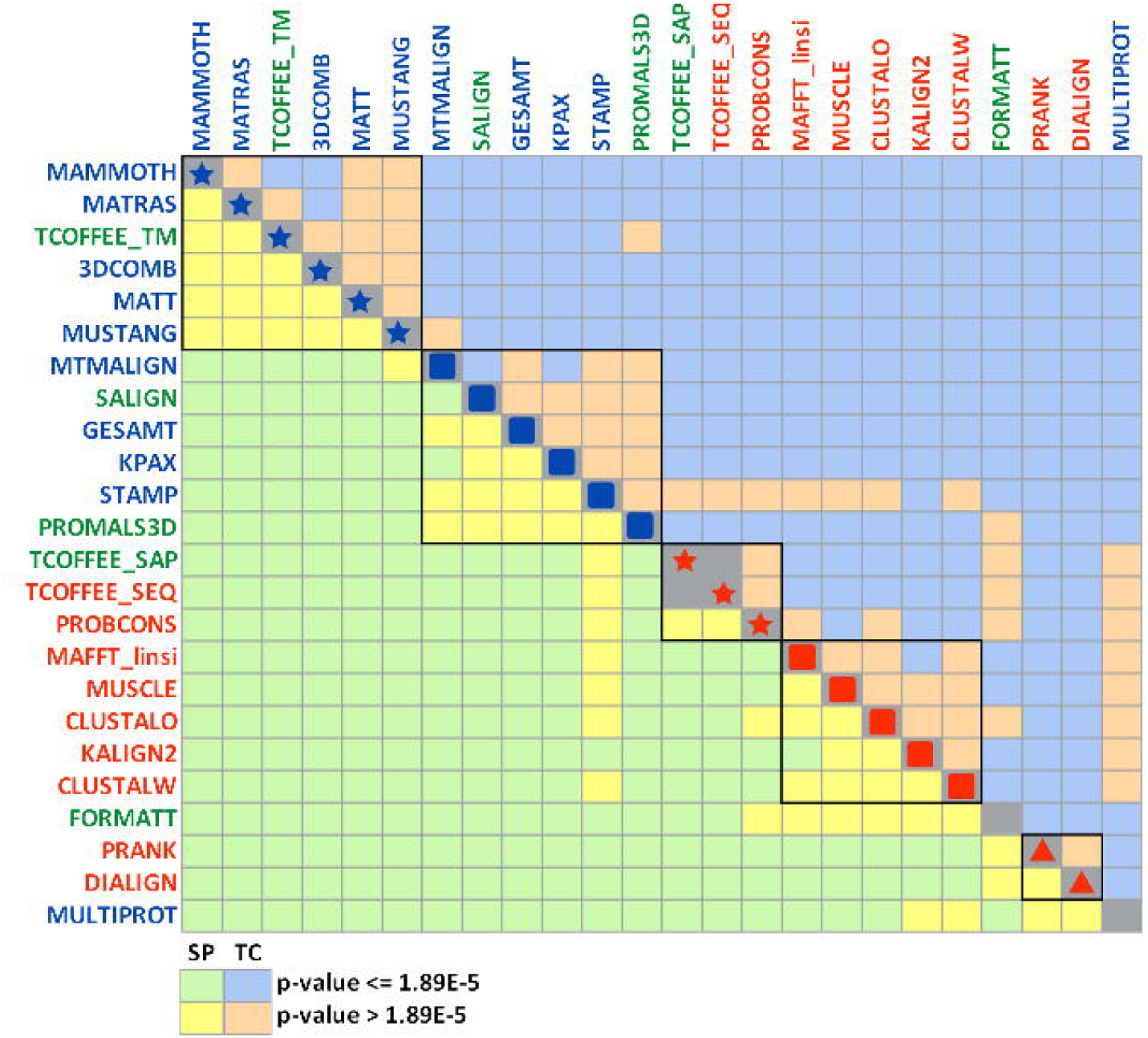
P-value heatmap of the Friedman tests Entries show the p-value computed using a Friedman rank test. Values above the diagonal where calculated with CS scores and values under the diagonal where calculated with SP scores. The programs are ordered according to their median SP scores. The colors of the boxes are: red for sequence-based alignment programs, blue for structure-based alignment programs, green for sequence+structure based alignment programs. The yellow or orange cells denote a non-significant p-value according to the 0.05 alpha risk, with a Bonferroni correction for multiple tests. The green or blue cells denote a significant p-value.

From this analysis, we can conclude that there is a ranking of groups of programs according to their overall performance, and that structure-based programs achieve better scores.

We also proceeded to hierarchical clustering on the basis of the scores of the various programs and the various alignments. A heatmap of this clustering is presented in Figure 4 for CS scores and in Supplementary Data Figure 1 for SP scores. The results are extremely similar with both scores. Considering alignment clustering, alignments for which all categories of methods succeed are more concentrated on the right side of the heatmap and the alignment clustering tree. In a thin central strip, the sequence-based methods have better scores: the score cells are in darker red for the structure-based. On the left side, the scores are better for structure-based and structure+sequence-based methods and in the extreme left side, all methods fail. Those difficult alignments are mostly from BB2, BB3 and SISYPHUS.

**Figure 4:**
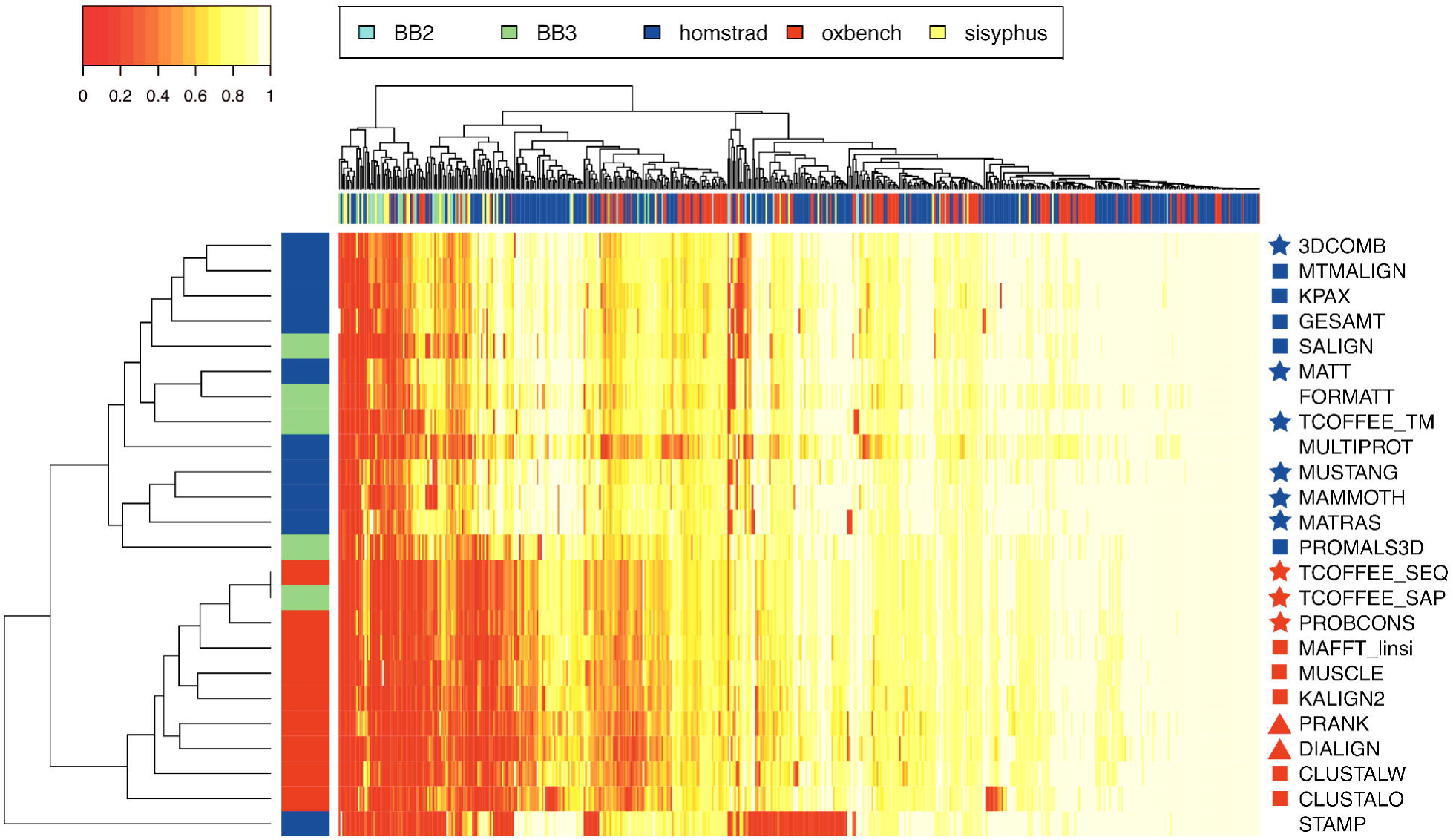
CS scores Heatmap and hierarchical classification of the programs and of the alignments (Complete method, Euclidian distance). The program colors are: red for sequence-based programs, blue for structure-based programs and green for sequence+ structure based programs. The symbols characterizing each program are the same as in Figure 3.

Considering program clustering, all structure-based programs but STAMP and sequence+structure-based programs but TCOFFEE_SAP are in the same sub-tree. All sequence-based are also pooled together. MUSTANG, MAMMOTH and MATRAS, which performances are undistinguishable according to the Friedman test, are very close in the tree. It means that their performances are similar, even if there are some discrepancies for some alignments. The next program in this branch is PROMALS3D, but its performances are significantly different from the three previous programs according to the Friedman tests. MATT and FORMATT are clustered together, but we know from the test that MATT results are better. The next closest method is TCOFFEE_TM. 3DCOMB, MTMALIGN, GESAMT and SALIGN are clustered together but according to the previous tests, 3DCOMB achieves the best results in this group of programs.

### The effect of sequence identity

We have investigated the effect of sequence conservation on the quality of the alignments computed by the different programs. The results are presented in Figure 5 for CS scores and in Supplementary Figure 2 for SP scores. As expected, the differences between structure-based and sequence-based methods are stronger for alignments of very divergent proteins. The difference is stronger in the case of CS scores but the effect is globally the same. It is more surprising to see that even at very high identity levels, structure-based programs still provide better scores than sequence-based programs. We also checked the effect of the number of proteins to align. The effect is very weak in the case of SP scores for all programs except MULTIPROT (see Supplementary Figure 3) but it is visible on the CS scores (Supplementary Figure 4).

**Figure 5:**
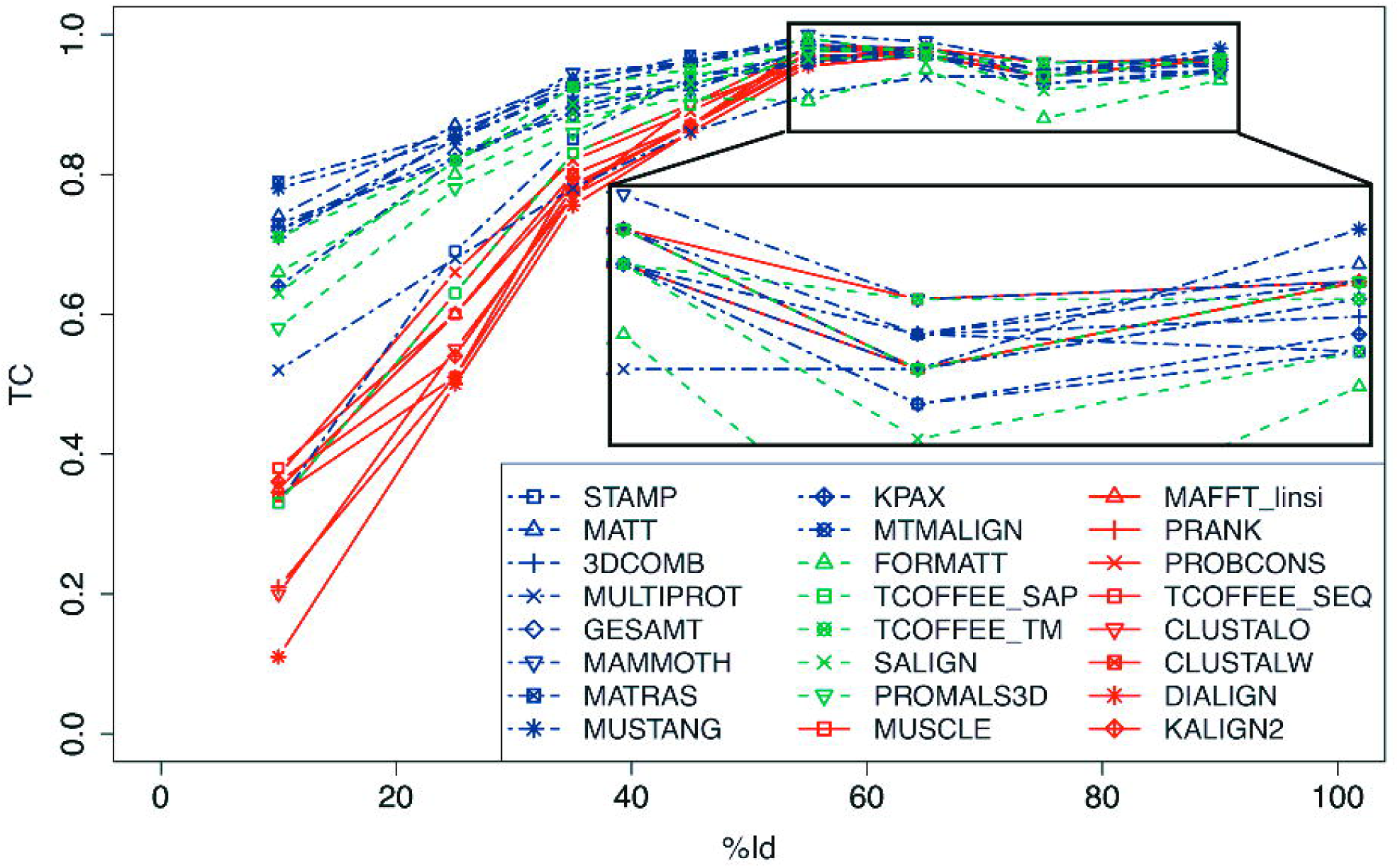
CS scores as a function of percentage identity of the core reference alignments

### SSE and burying effect

We also investigated whether structure-based methods are strongly dependent on secondary structures and solvent exposure. We computed the SP and CS scores independently core residues in helices, strands or loops; the same procedure was applied for exposed or buried residues. The results are presented Figure 6 for CS scores. The scores decrease for loop residues; this decrease is more importantly for structure-based and structure+sequence based methods than for sequence-based methods. Similarly, the scores decrease for exposed residues for all methods. In summary, buried or regular secondary structure regions are better aligned by all programs than exposed regions or loops.

**Figure 6:**
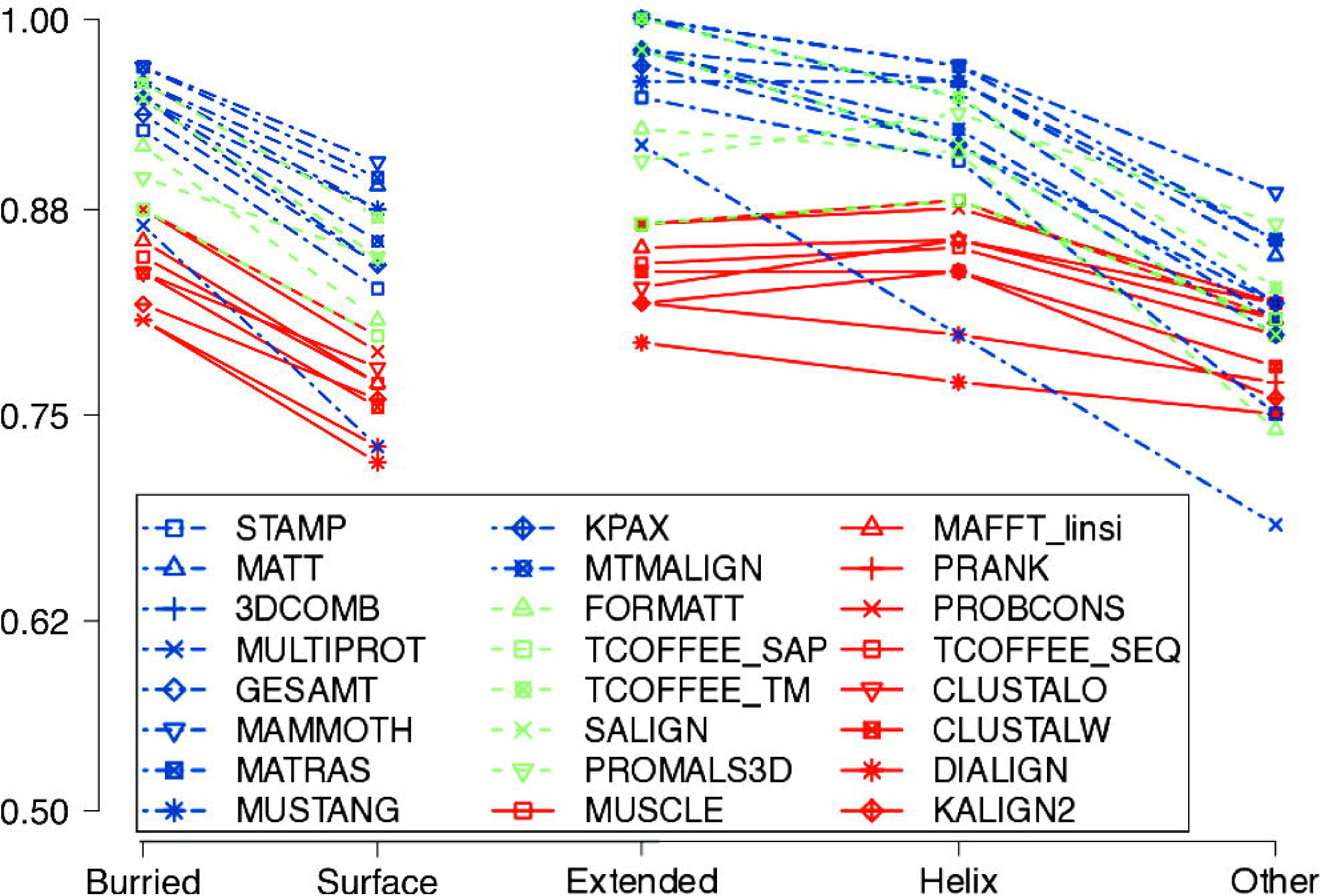
Median CS ccores for each program. Left: core columns where the residues are either in helix or strand or loop. Right: residues either buried or exposed.

### Database effect

We wondered if the success rate of the programs was dependent on the databases. The composition of the various databases is different in terms of sequence identity and core definition. We tried to remove these this biases by selecting alignments between 10% and 40% of sequence identity, because all databases are present in this range. Besides, only core positions in conserved regular secondary structures were selected. In Figure 7, it is clear that the CS scores fluctuate depending on the reference alignment origin: the median scores are globally higher and less variable for the two HOMSTRAD and OXBENCH which contain more alignments and whose generation procedure is more automatic than BB2, BB3 and SISYPHUS. However, the ranking of the programs is similar: the same structure-based or structure+sequence-based programs are the best, even if their order slightly varies. The most affected program is STAMP, which performances are poorer with the three last databases.

**Figure 7:**
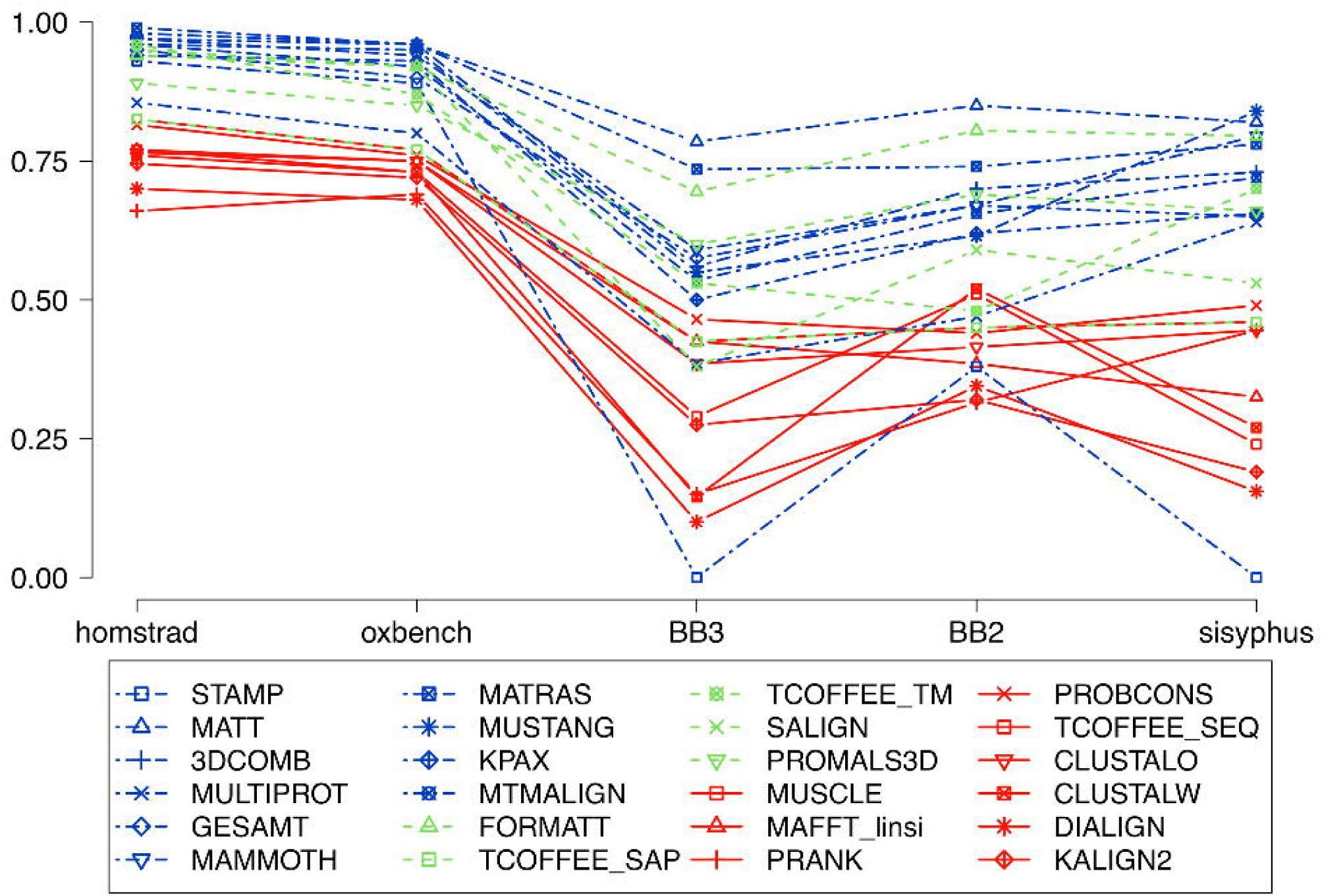
Median CS scores for each program and each database, restricted to alignments in the range 10-40% sequence identity. Besides, only core positions in regular secondary structures perfectly conserved in all the proteins of the family are considered. Color code is the same as in Figure 6.

### RMS and sequence identity

The multiple RMS among proteins of the families are smaller for structure-based methods than for sequence-based methods as expected because structure-based methods align proteins while optimizing the structural resemblances (see Figure 8). The RMS computed according to the reference alignments (Figure 8, black line) are in between the two categories except for alignments above 70% identity where these RMS are higher. These high RMS are mostly from OXBENCH alignments and are mainly due to an alignment in multidomain chains. The best programs, as resulting from the previous sorting, are not those with the smallest RMS; on the contrary, the order is globally reversed. Figure 8 highlights the differences in the goal of the programs: either optimize structural similarities or consider some evolutionary aspects. As we are testing the capabilities of programs to retrieve homologous positions, the second category of programs is advantaged in our study: if they are better here, it is only in their capacity at retrieving homologous positions, but they may miss other structural similarities.

**Figure 8:**
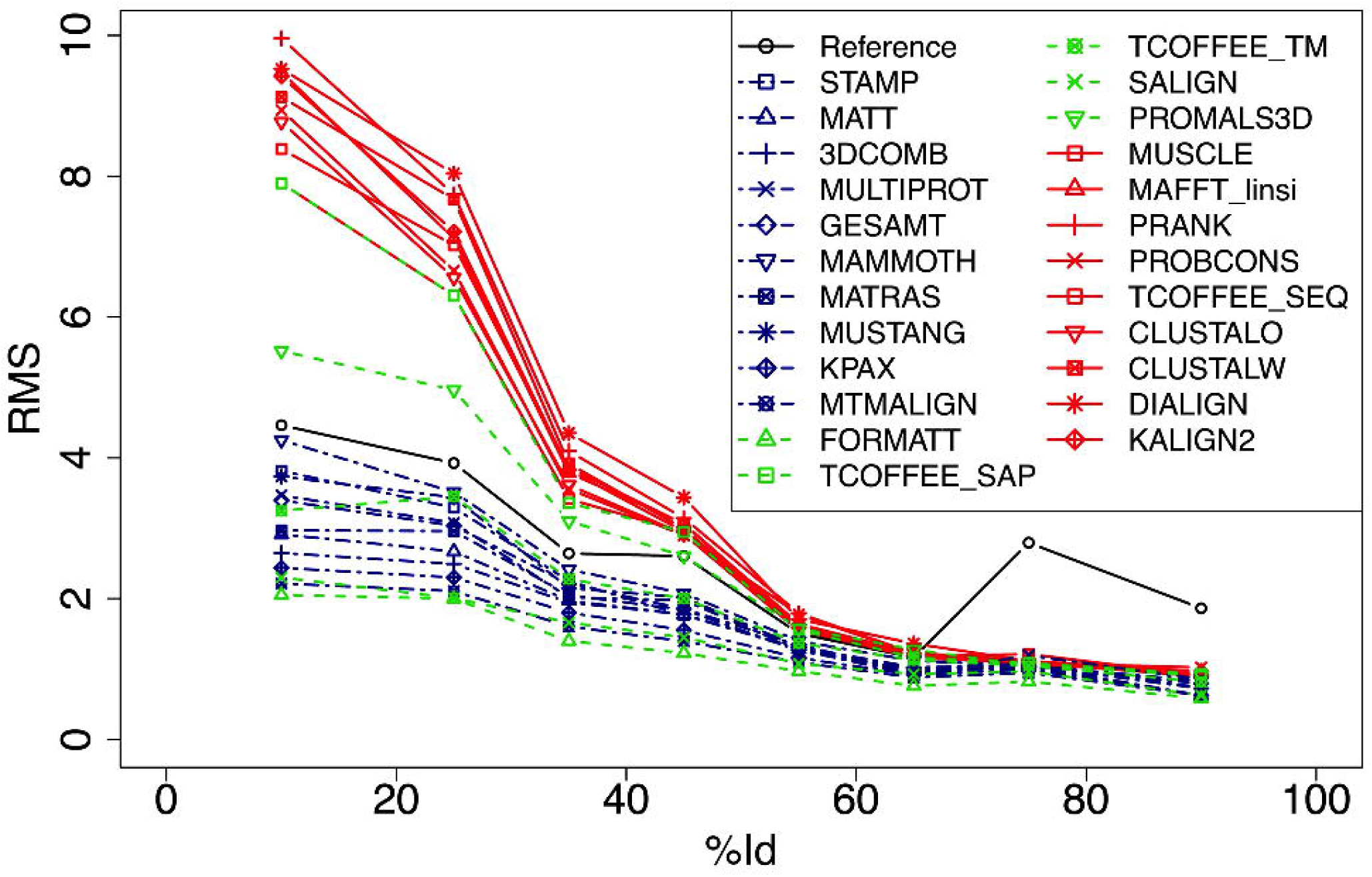
RMS as a function of core alignments identity percentage

### GAPS

The proportion of gap opening is clearly different in sequence-based and structure-based programs (see Figure 9). The structure-based programs but MAMMOTH tend to over-estimate the number of indels and the sequence-based tend to under estimate the number of gaps. MAMMOTH has a linear penalty gap function which seems to be quite efficient. PROMALSD3D has also a linear gap penalty function and tends to place less gaps than in the reference alignments. PRANK, which has been designed to correctly place the indels, is the closest method to the reference. As most of the structure-based methods work with small structural blocks, they don’t have a gap penalty function, which explains this possible over-estimation. We believe that some improvement in the gap treatment for structure-based and sequence+structure based methods should improve their performance.

**Figure 9:**
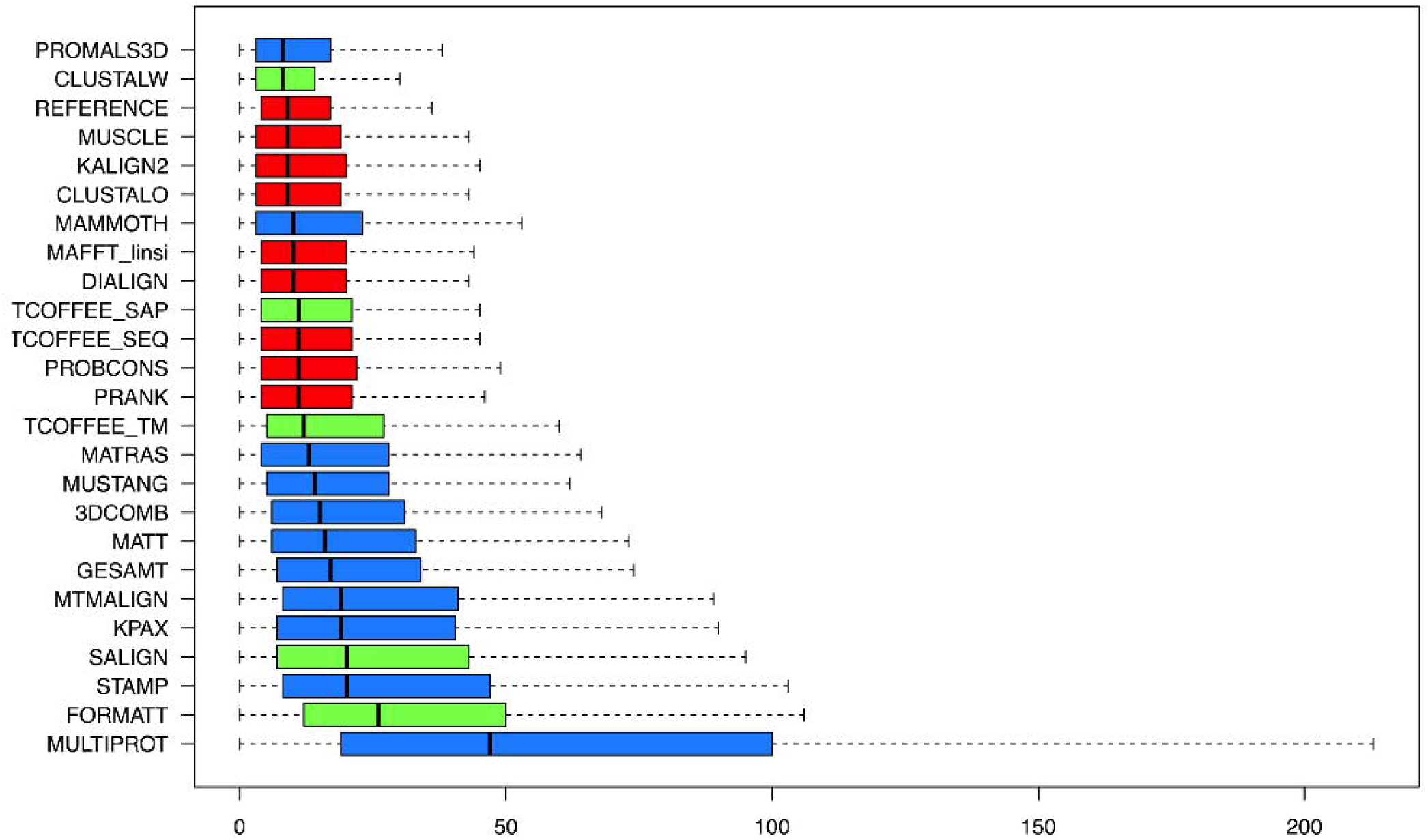
Distribution of number of alignment columns containing one or more gap opening. The colors of the boxes are: red for sequence-based alignment programs, blue for structure-based alignment programs, green for sequence+structure based alignment programs, grey for reference alignments.

## DISCUSSION

In this article, we have compared the ability of sequence-based, structure-based and sequence+structure based multiple alignment programs to retrieve homologous positions of defined in reference alignments from five well known datasets. The structure-based programs have globally better performances than sequence-based, but also than most of the structure+sequence-based programs. A group of five structure-based plus one sequence+structure-based programs are significantly better than the others: MAMMOTH, MATRAS, TCOFFEE_TM (sequence+structure), 3DCOMB, MATT and MUSTANG. All those six programs build the alignments from pairwise aligned fragments of few residues. No obvious superior methodology has been identified. However, it is interesting to notice that those program performances are different, according to a hierarchical clustering of their scores: they do not all cluster together, meaning that their success or failure depends upon the alignment. It is therefore possible that a consensus method achieves better results if it is possible to identify the cases where each method succeeds, as it have been suggested also in the article of Berbalk et al. (25). The performance differences between sequence and structure-based programs are stronger for low identity alignments as it has been highlighted in Kim and Lee (23), but they are still present at high identity. They weakly depend on the 3D localization of the residues.

TCOFFEE_TM is the only sequence+structure-based program in the group of best programs. The adjunction of structure information clearly improves the alignment done by TCOFFEE_SEQ. In sequence-based programs, in our study, the consistency-based programs (TCOFFEE, PROBCONS) are the best ranked as in (14, 51) but without MAFFT. These consistency-based methods are quite efficient and it would be interesting to see their results with only structure information. We can conclude that while aligning proteins for the identification of homologous positions and if it all structures are known, it is better to align the protein with their structures. In the case where not all structures are known, it is probable that it is better to use a sequence+structure-based method as TCOFFEE_TM, but this exact case has not been addressed in this study. Knowing the difficulty of combining structure and sequence information, this case has to be tested before counseling it.

The second performance group of structure-based programs contains four structure-based programs (MTMALIGN, GESAMT, KPAX and STAMP) and two sequence+structure-based programs (SALIGN and PROMALS3D). STAMP is apart, but all three other structure-based programs tend to produce low RMS alignments; they are also clustered together. These programs seem to be more dedicated to the identification of structurally similar regions, which are not always homologous. It would be very interesting to compare the regions identified as structurally similar by those programs and not by the six previous ones: these regions may sequence similar-structure dissimilar which would help to understand the complex evolution of protein structure. STAMP has a very different behavior which renders comparison difficult: globally, it succeeds for databases that are more automatically constructed (HOMSTRAD and OXBENCH) and mostly fails for the others (BB2, BB3 and SISYPHUS). However, it is surprising that its performances are not the best for the two first databases as it is used to compute their alignments. Maybe the program version or parameters are different, or the afterwards refinement of the reference alignments may explain it. MULTIPROT is also very different from the other structure-based programs: it is dedicated to the identification of reliable aligned columns which are locally structurally similar. It does not align the other regions, which explains its poor scores. However, it is very efficient for column identification: its CS score are clearly better than its SP scores, and it is the only structure-based program with this behavior.

The scores of all programs and their dispersions are similar for the two databases HOMSTRAD and OXBENCH which are the most automatically generated. The scores are different for the three other databases which are more manually built: they are globally lower and more variable among the programs, meaning that these alignments seem to be more difficult to retrieve. Whatever the database used, the first ranked program is always a structure-based program. Although, structure-based and sequence+structure-based programs have better scores than sequence-based programs. However, the ranking may vary: in the subset of alignments from the three manual databases, with 10 to 40% of identity and only the positions perfectly conserved in terms of SSE, MATT, MATRAS, FORMATT and MUSTANG achieve the best results. Another bias in this study is that these 5 benchmarks are built from protein structural information which may advantage structure-based methods. Other broad of benchmarks exit (12). It is not possible to use simulated sequences in the case of structure-based programs but it would be interesting to compare the program alignments altogether without a reference to check their consistency. The last type of benchmark is to compute phylogenetic trees from the program alignments to compute a score from the correctness of the trees. This analysis could also be done for all type of programs.

All structure-based programs except MAMMOTH and all sequence+structure-based programs except PROMALS3D and TCOFFEE_SAP have a greater proportion of columns with a gap opening than reference alignments and all other methods have a lesser proportion. Most of structural-based methods don’t have a penalization function of gaps which explains this behavior. It is possible that a penalization of gaps would improve the alignment quality.

Finally, some improvement improvements concerning usability and applicability of structure-based programs would generally been valuable.

## CONCLUSION

We can conclude from this study that it is indeed better to use structure information than sequence information only to identify homology in proteins, but the difficulty of combining sequence and structure information is obvious: the sequence+structure-based methods are not better than the structure-based method. Several programs are globally equivalent in performance but their behavior vary for each alignment and maybe, a consensus method could achieve better results. However, a real model of sequence and structure protein evolution would surely greatly improve the methods but such a model is quite difficult to design notably because of the folding process which may drastically change the structure even if the sequence difference is not that strong. There is also still room for improvement in term of software ergonomics and gap treatments.

## AVAILABILITY

All results presented in this study are available upon request to the corresponding author.

## ACKNOWLEDGEMENT

The authors greatly appreciated the informal discussion with the authors of most of the programs used in this study.

## Abbreviations

ASA: Accessible Surface Area
CDD: Conserved Domain Database
MSA: Multiple Sequence Alignment
PDB: Protein Data Bank
RMS: Root Mean Square
SP: sum of pairs
CS: column score
DP: Dynamic Programming
AFP: Aligned Fragment Pairs
SSE: secondary structure element

